# Variability of wheel running behavior in mice is dependent on housing, sex, and genetic background

**DOI:** 10.1101/2022.10.28.514233

**Authors:** Zara Kanold-Tso, Dietmar Plenz

**Affiliations:** Section on Critical Brain Dynamics, National Institute of Mental Health, Bethesda, MD, 20892, USA

**Keywords:** Running, transgenic mouse, locomotion, behavioral phenotype, competition

## Abstract

Environmental enrichment (EE) has been increasingly used to enhance and support the physiological fitness and natural behavior of research animals. For mice, which are known to be highly active nocturnal animals that live in large social groups, the introduction of running wheels is a particularly promising approach in EE. When wheels are introduced to their home cage, mice increase their nocturnal locomotion activity by orders of magnitude. However, little is known about how mice share such a readily adopted EE. Here, we studied wheel running in single-housed and group-housed mice. We hypothesized that group-housed mice will compete over the scarce resource of a running wheel leading to a change in overall wheel usage compared to single-housed mice that have exclusive wheel access. We measured wheel locomotion in two strains of single-housed and group-housed mice over multiple 24h periods at sub-second temporal resolution using a custom-designed data acquisition system. We observed both sex- and housing-specific differences in wheel usage. In single-housed C57Bl/6 mice, mice ran at consistent speeds and females ran larger distances. In group housing, periods of slow locomotion speeds as well as speed transitions emerged with fewer breaks in wheel usage. This group effect was less pronounced in a genetic mouse model (parvalbumin-Cre on C57Bl/6 background).

Our results demonstrate a change in wheel usage during group housing which supports competition over EE resources while enhancing overall locomotion in mice.

## Introduction

Animal research is fundamental to science. Accordingly, the well-being of animals is essential to research and development and is commonly supported by environmental enrichment (EE) during animal breeding and animal housing. For mice, social housing, nest building material, and hiding places such as narrow tubes are now commonly used components of EE. In contrast, EE that supports the natural tendency of mice to run long distances is more challenging to implement within the constraints of research facilities. To accommodate such long-distance running, running wheels can be introduced into the home cage. The running behavior of mice has been studied by placing running wheels inside cages and instrumenting running wheels with sensors, capturing the numbers of wheel revolutions. Such studies have shown that mice run great distances of up to 20km/day (De Bono et al., 2006; Goh & Ladiges, 2015; Koteja, Garland, et al., 1999; Koteja, Swallow, et al., 1999; Zhu et al., 2021) and that most running episodes occur during the dark cycle (Fuochi et al., 2021; Kobayashi et al., 2020). Moreover, mice tend to run at a characteristic “cruising speed” (De Bono et al., 2006). On the other hand, EE resources might cause conflict between mice. For example, mice are often group-housed to provide for social enrichment and the introduction of a running wheel into group-housed cages could potentially set up a competitive situation for the running wheel, which might alter running behavior. Here, we address this potential conflict by comparing running wheel activity in single- and group-housed mice.

Transgenic mice are commonly used in neuroscientific studies to study the effects of specific genes or to manipulate cells based on the expression of specific genes. One commonly used technology for expressing genes in a selected group of cells is the expression of Cre recombinase in specific cells. Mice carrying Cre recombinase are then often crossed with mice carrying genes that are flanked by loxP sites (floxed mice) (Kim et al., 2018). Conditionally genetically manipulated mice are then compared to unmanipulated wild-type mice. The assumption is that the presence of Cre-recombinase by itself does not have a physiological effect. Parvalbumin-expressing neurons are a major group of inhibitory interneurons in the brain (Hu et al., 2014), and parvalbumin-Cre (PV-Cre) mice (JAX #008069 on C57Bl/6 background) are commonly used to manipulate inhibitory interneurons to infer the effect such neurons have on neural activity and behavior by comparing to unmanipulated wild-type mice of the same strain, e.g. C57Bl/6. The assumption in such studies is that the expression of Cre-recombinase does not influence neural activity or behavior. We thus tested if the expression of Cre by itself will alter running behavior by comparing the running behavior of Parvalbumin-Cre mice with C57Bl/6 mice.

We instrumented a running wheel and measured detailed running behavior by measuring the speed of running episodes and the changes in speed between episodes. We find that in C57Bl/6 mice group housing leads to a larger variability in running speeds and that this difference between single- and group-housing is most pronounced in female mice. We did not observe differences in running patterns between single and group-housing in PV-Cre mice.

## Results

To measure running in populations of mice over long periods of time we built an instrumented running wheel that could be placed into the home cage 24/7 (Fig. 1A). We measured wheel running in single-housed (n=18 cages, 9 female, 9 male) and group-housed C57Bl/6 animals (n=8 cages with n=3 animals each. 4 cages females, 4 cages males) over a period of 2 – 3 days each. During this time, the animals had free access to the wheel. The running wheel was instrumented with four magnets and thus allowed the collection of speed data with ¼ revolution precision. We measured wheel motion continuously at a rate of 100Hz and calculated instantaneous running speed.

**Figure 1:**
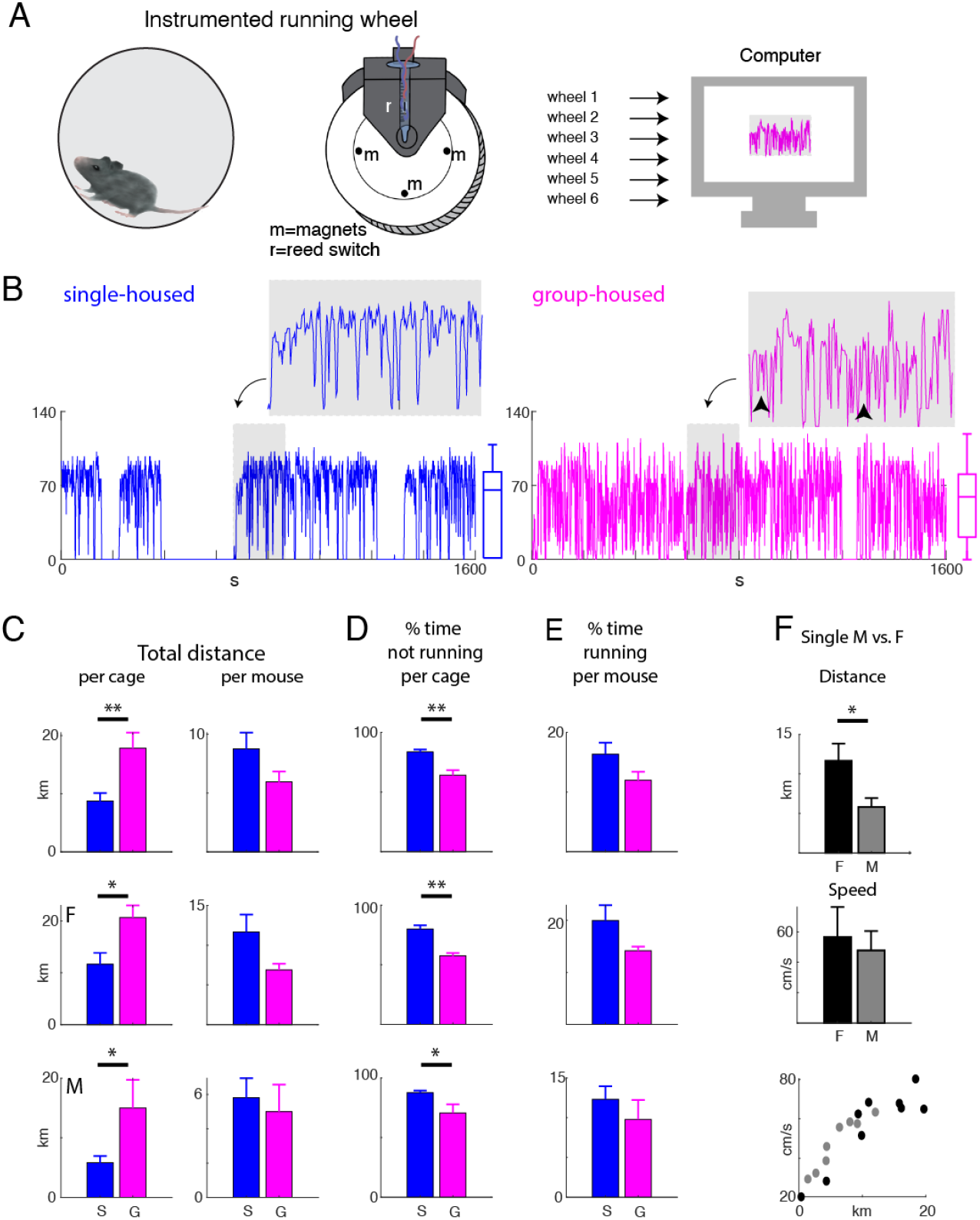
Instrumented running wheel to measure mouse locomotion. **A**: Experimental setup. The running wheel is instrumented with 4 magnets and a reed switch. Up to six wheels can be recorded simultaneously through using National Instruments NI board connected to a stand-alone computer. **B**: Shown are examples of instantaneous running speeds from a cage of single and group-housed mice. Running speeds in single-housed mice appear to switch between 0 and ~70cm/s, whereas for group-housed mice speeds are more diverse. **C**: Bar graphs showing the total distance covered in 24h per cage (left column) and per mouse (right column). Top row is all cages, the middle row is females, and the bottom row is males. Cages with group-housed mice show a larger total distance covered (p<0.05, Student’s T-test). **D**: Fraction of time mice in each cage did not run (p<0.05, Student’s T-test). **E**: Fraction of time running per mouse (p>0.05). **F**: Distance and speed for single-housed male and female mice (‘*’ p<0.05). Distance run grows with average running speed.

### Running distance is dependent on housing conditions

Mouse locomotion data was measured using the simple counts of wheel rotations converted into distance over time. Displayed in Fig. 1B is the running speed during a 1600 second interval of wheel usage during mouse locomotion for both a single- and group housed-mice (blue and magenta respectively). A zoomed in view into the timeframe for an 200 second period displays the speed patterns for both the single housed and group housed mice in further detail. Using box plots to display the bulk of running speeds over time, we can already see in two instances of single vs group housed mice that the bulk of the single housed mouse’s running was at a more consistent speed than the group-housed mice (Fig. 1B).

Group-housed mice covered up 20km over a 24hr period on the running wheel, with groups of mice running about 19km. Groups of males ran about 15km, while groups of females ran about 21km (Fig. 1C). In single-housed settings, the females ran about 11km in the 24hr recording period and males about 5km, averaging the single-housed mice to about 9km in 24hrs (Fig. 1C).

To investigate if group-housing changed the distance each mouse ran, we calculated the average distance travelled for each mouse. To estimate the distance of each mouse in the group cage covered we divided the group-housed distances by three. The average distance was about 6km in the 24hr period, with about 7km average for females and 5km for males (Fig. 1C). While we observed a trend for group-housed females to run less than single housed females, this was not significant due to large variability (p>0.05). Assuming each mouse in the group setting ran exactly 1/3 of the total group distance, this would mean that group-housed mice run on average similar distances than their single-housed counterparts.

Competition or hogging of the wheel may influence average running distance or running time for each group member, compared to the single-housed mice. In order to further examine the dynamics of when single and group-housed mice are using the wheel, we analyzed the time spent running and not running for both housing situations.

In Figure 1D it is demonstrated that single-housed mice, both male and female, are inactive for a longer period of time than the total male and female group-housed mice (3 mice per cage). When the inverse data is taken, so period of time spent running, and then divided by three to determine average mouse running time per group-housed mouse, the single-housed mice prove to be active for a similar period of time that each group-housed mouse (Fig. 1E). In correspondence with the trend to lower running distances in females in Fig. 1C, we here also observe a trend to lower wheel utilization in group-housed females than single-housed female mice.

Though there are more mice in the group cages, they do not prove to run for longer time nor distance than single-housed mice. This may be due to inept ability to use the wheel for long enough periods of time to rack up distance, due to increased competition over the wheel or dissuasive group dynamics causing wheel use to be overall discouraged by some (possible alpha) mice. Further inspection through video or RFID monitoring would be needed to determine the true behavioral dynamics causing the changes in wheel use and true group dynamics.

Comparing single-housed males with single-housed females showed that females covered larger distances than males (p<0.05, T-test) (Fig. 1F top). Calculating the average speed when running showed that while males and females ran at similar speeds (Fig. 1F middle), speed and distance were correlated in that slower males and females ran shorter distances (Fig. 1F bottom). These data indicate that the large variability seen in running distances might be due to differences in the running speeds of individual mice.

### The distribution of running speeds depends on housing conditions

To further examine the probability of different running speeds in group vs. single-housed mice we looked at the fractions of time spend at each possible speed for group- and single-housed mice. The probability of instantaneous running speeds of 3 randomly selected cages for both female and male, group and single-housed mice are shown in Fig. 2A and Fig. 2B. Both male and female single-housed mice display one or two peaks in the running speed distribution, meaning they most consistently run at either of these speeds (Fig. 2C). Male and female group-housed mice on the other hand contain on average two to three peaks in speed (Fig. 2D). Quantifying the numbers of peaks in the speed distributions for each cage shows that single housed mice show a majority of one peak in running, and rarely more than one, whereas group housed mice often show more than one speed peak (Fig. 2C, D) (p<0.05).

**Figure 2:**
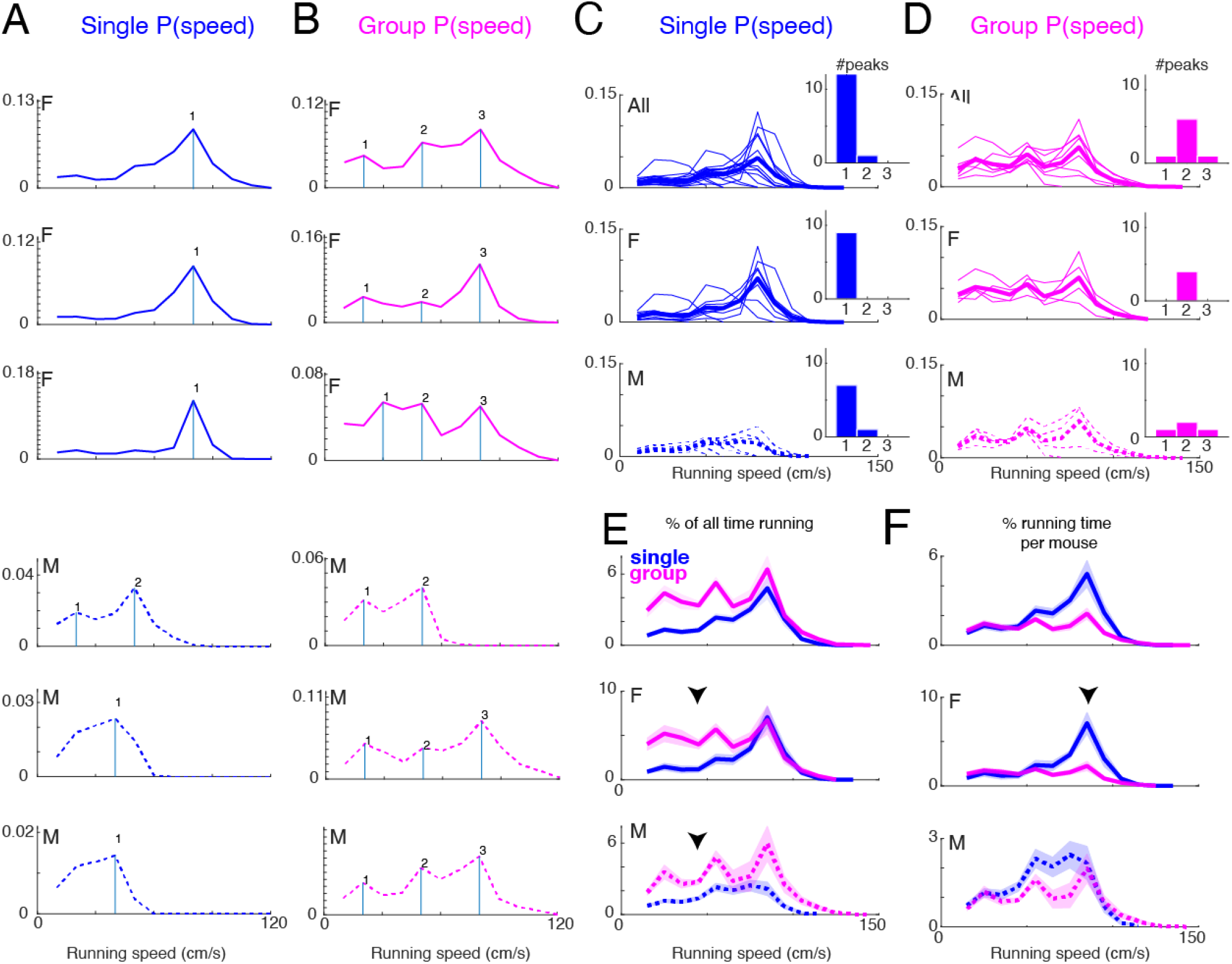
Speed probability distributions of group-housed cages show multiple peaks. **A, B**: Plotted is the distribution of instantaneous running speeds over the recording period for 6 example cages of single-housed (A) and group-housed (B) mice. Vertical lines and numbers indicate detected peaks in the distributions. Distributions from females in solid lines, males in dashed lines. **C, D**: Superposition of traces in A and B (thin lines), respectively, and the average trace of all single- or group-housed cages (thick line). Insets show the distribution of the numbers of speed peaks. Most single-housed cages showed only single peaks, while group-housed cages had multiple peaks (all p<0.05, KS test). **E**: Comparison of mean (±SEM) speed distribution for single and group-housed cages. Note that single-housed cages show a prominent single peak in the distribution. Group housed mice show more running at low speeds (arrowheads). **F**: Comparison of mean (±SEM) speed distribution for individual mice from single and group-housed cages. Note that single-housed mice but not group-housed mice show a prominent single peak in the distribution (arrowhead).

When the average speed of both single and group house mice are graphed together, the definitive trend is established that group-housed mice contain more running speed peaks (2-3) than single housed mice (1-2) (Fig. 2E). Group-housed mice contain the same mutual peak at a high speed (~80cm/s) as single-housed mice, but also contain one to two additional peaks at lower speeds than single-housed mice (arrowheads), meaning group-housed mice are more probable to spend more time running at the lower speeds than single-housed mice (Fig. 2E).

Out of all time that the mice are running (speed> 0), both male and female group-housed mice run on average more of the time at lower relative speeds than single-housed mice (Fig. 2E). When time running was divided by 3 for group cages, to examine individual running time at each speed per mouse, group-housed mice female mice individually ran less percentage of the time at high speeds (~80cm/s) than single housed mice (Fig. 2F, arrowhead).

In group-housed conditions, male and female mice show no obvious pattern differences between probability of running speeds over time, but single-housed female mice show a more pronounced peak at higher running speed (Fig. 2F, arrowhead) than male mice.

This shows that both male and female group-housed mice spend more time running at lower speeds compared to single housed mice and that this effect is more pronounced in female mice. This change in running speed is consistent with the trend to reduced running distances in group-housed female mice (Fig. 1C). This effect of group-housing might be due to increased competition about using the wheel or transitions where mice get in and out of the wheel (where they will therefore not be moving at a high speed).

### Group housed mice show more variability in running speed transitions

So far, our results have shown that group-housed mice spend more time running at lower speeds. Moreover, running traces of group-housed mice seemed to show more speed transitions (Fig. 1B). To examine the nature of these speed transitions quantitatively, we computed the speed changes occurring by plotting the running speeds between time bins during running intervals. Instantaneous running speed traces were evaluated at times n and n+dt seconds (for dt=1, 5, 10, and 100s delays) to determine the transition probability in the mice locomotion speed (Fig. 3A). The left example shows an example from a single-housed cage. The two red lines in the trace show the times the running speed was evaluated. The two speeds were similar. In contrast, the trace on the right was obtained from a group cage. The running speeds in the evaluation interval were different. To assess the overall pattern of speed change in each cage we evaluated running speed transitions for all time bins and plot calculated the probability particular speed pairs, speed(tn) and speed (tn+1) occurred in a recording. Figure 3 (B, C) shows example plots of the probability of speed transitions in single and group-housed cages with female and male mice. Bright warm colors indicate a high frequency of speed pair occurrences (dark blue indicates no occurrences). For single-housed mice, we typically observed a single tightly circumscribed area indicative of stable running spread, e.g., the speed at time tn and at time tn+1, was similar (Fig. 3B, C). However, for group-housed mice, the speed areas were spread out over a larger area and could show multiple discrete areas indicating distinct combinations of speeds that were common. This suggested that mice under group-housed conditions showed many different variations in running speed but that certain speed combinations were more common.

**Figure 3:**
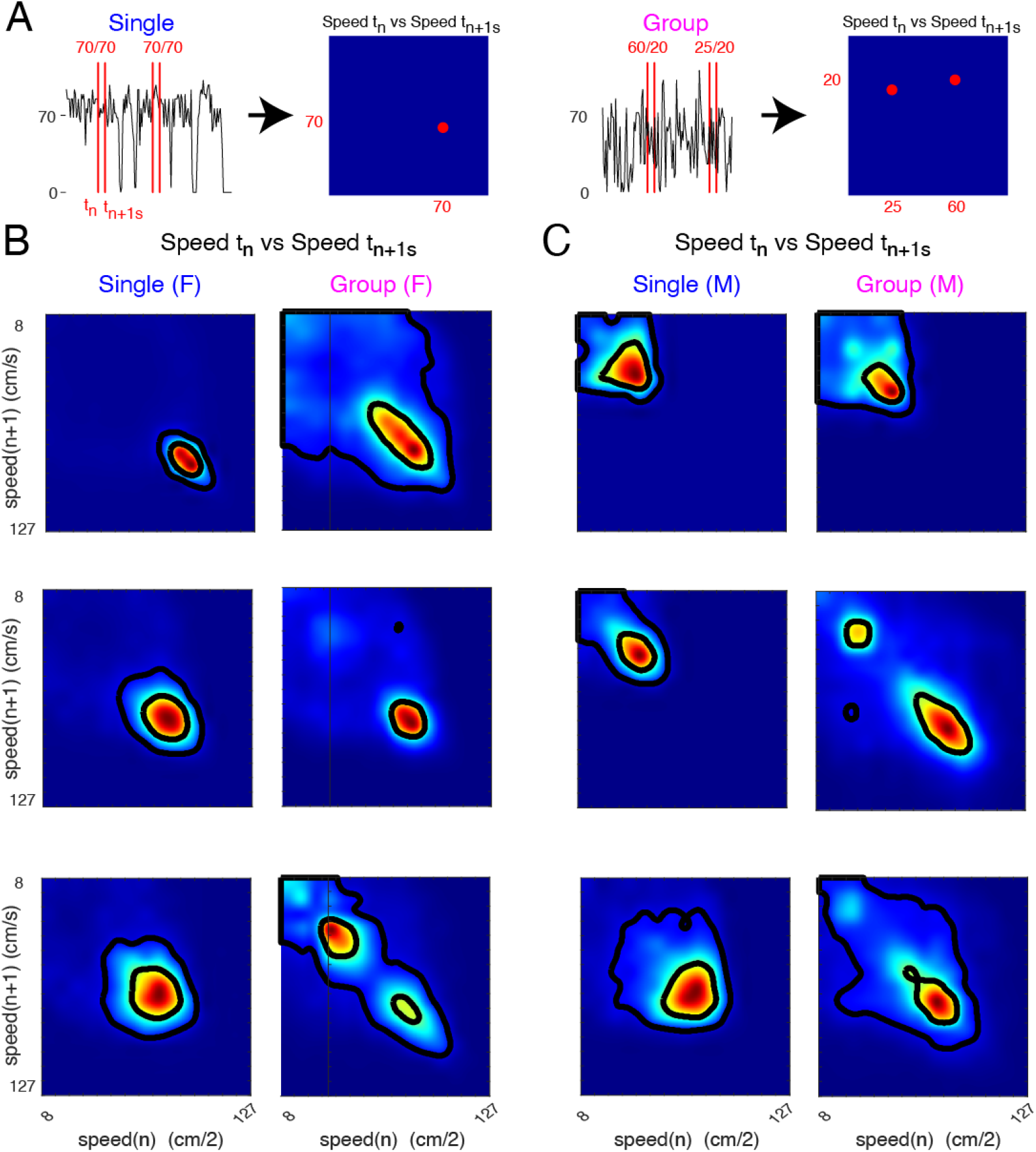
Transitions in running speed are more variable in group-housed mice. **A:** Schematic illustrating calculation of running speed transition. Speed is measured at time n and at time n+dt (dt = 1s; vertical red lines). The left trace is from a single-housed cage, and the right trace is from a group-housed cage. Note that speeds are more variable in group-housed cages. A histogram of each pair of measured speeds for all times n is then computed. **B, C**: Histograms of n/n+1 speeds for 3 cages each of single-housed and group-housed female (B) and male (C) mice. The color indicates the probability of each particular speed pair. Note that in single-housed mice, speed pairs form single circumscribed areas, while in group-housed cages, multiple areas are visible, and areas seem broader. Black lines indicate area boundaries at a low (10%) and high (50%) threshold.

To quantify these qualitative observations, we first counted the number of distinct speed transition areas for different delays (e.g. 1s, 5s, 10s, 100s) (Fig. 4). We found that for delays of 1s and 100s group housed cages showed more speed areas than single housed cages. This indicates that mice in group-housed cages showed more distinct patterns of speed transitions. We next quantified the compactness of the speed areas by measuring the size of the 2-dimensional probability distribution at two different thresholds (10% and 50%) and then computing the ratio (R=Area50%/Area10%) (Fig. 5A). A compact distribution would be expected to have a similar size and this a ratio close to 1, while a more distributed distribution would have a higher ratio. We found that for all delays, the area ratio was higher in group-housed cages than in single-housed cages (Fig. 5B-E). Moreover, this effect was present in both male and female mice, except that male mice did not show a significant difference at the longest delays. To shed more light on the origin of the change in ratio, we plotted the areas for the different thresholds (Fig. 6A, B) and find that the speed-transition area for the lower threshold was larger in group-housed cages than in single-housed cages (Fig. 6A), while the high-threshold area was similar (Fig. 6B). This indicates that group-house mice have a more variable running pattern in that they show a larger range of speed transitions.

**Figure 4:**
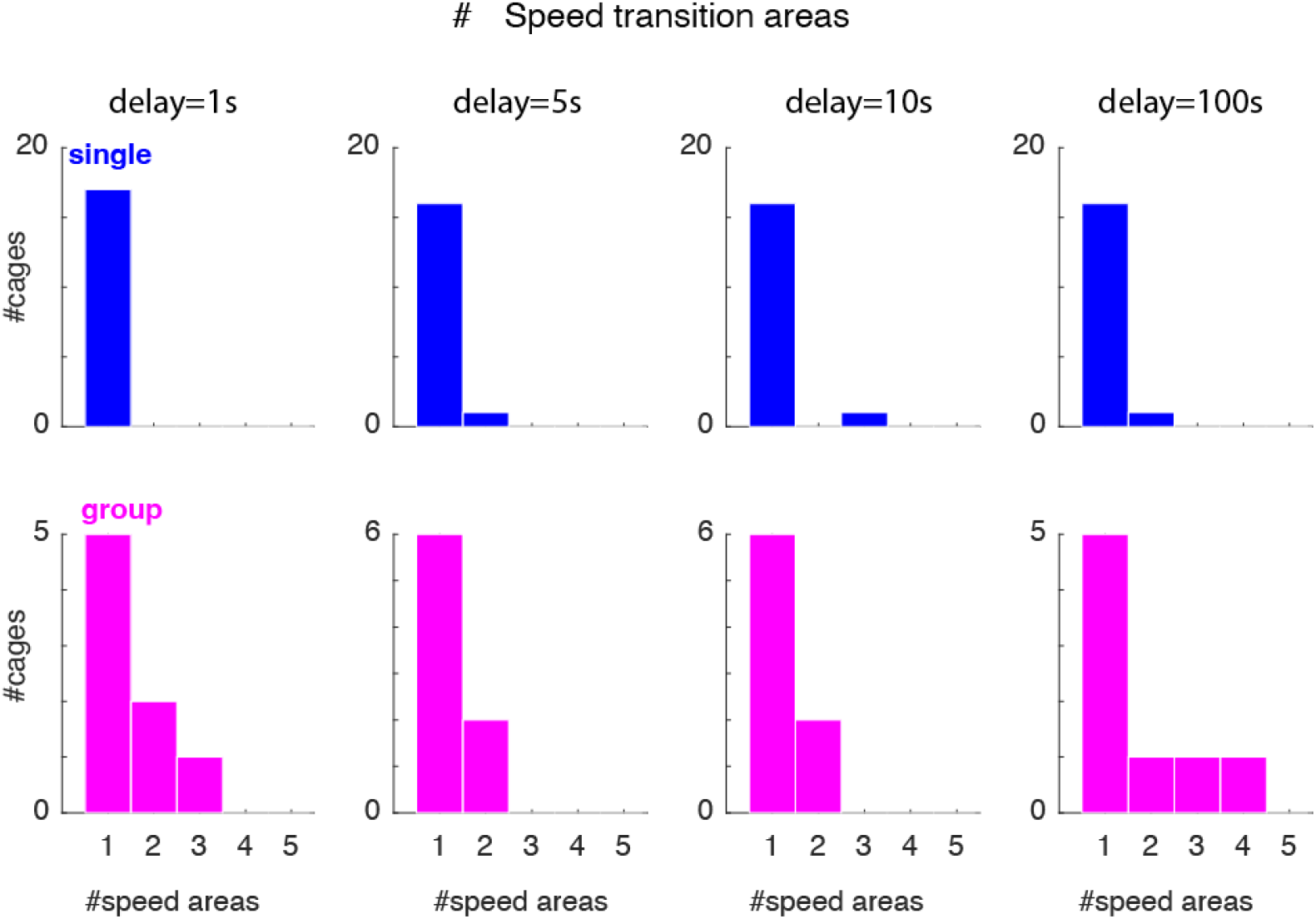
Group-housed mice have more speed transition areas. Histograms show the number of speed transition areas in single- and group-housed mice for delays of 1s, 5s, 10s, 100s. Group-housed cages show more speed transition areas for delays of 1s and 100s (p<0.05 KS-test).

**Figure 5:**
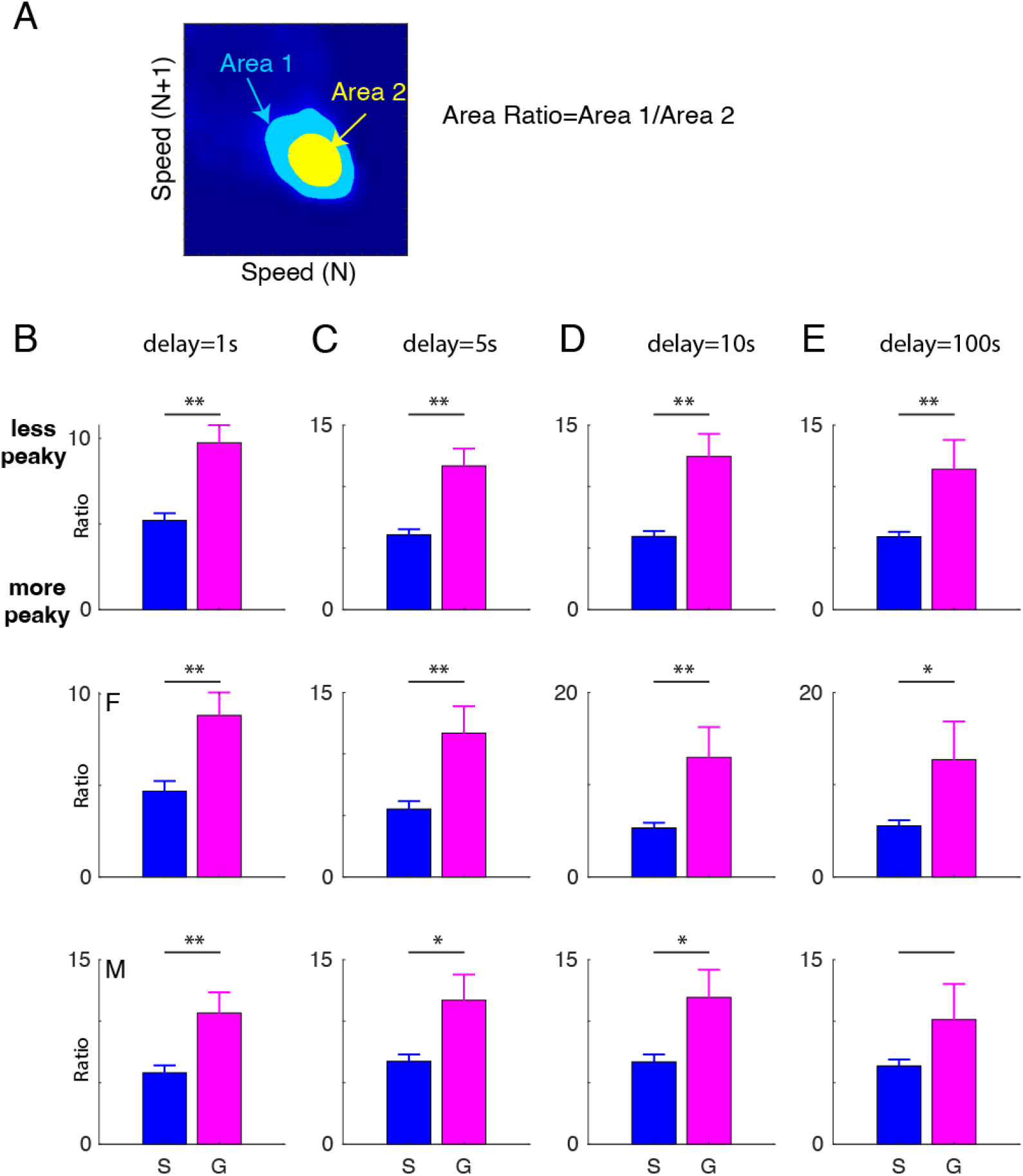
Group-housed mice have more peaked speed transition areas. **A**: Schematic showing how peak index is computed. Areas are calculated with boundaries at a low (10%), and high (50%) threshold, and then the ratio of the areas is computed. Histograms show the peak index in single- and group-housed mice for delays of dt = 1s (**B**), 5s (**C**), 10s (**D**), 100s (**E**). Group-housed mice show a higher peak index for almost all conditions (p<0.05 T-test).

**Figure 6:**
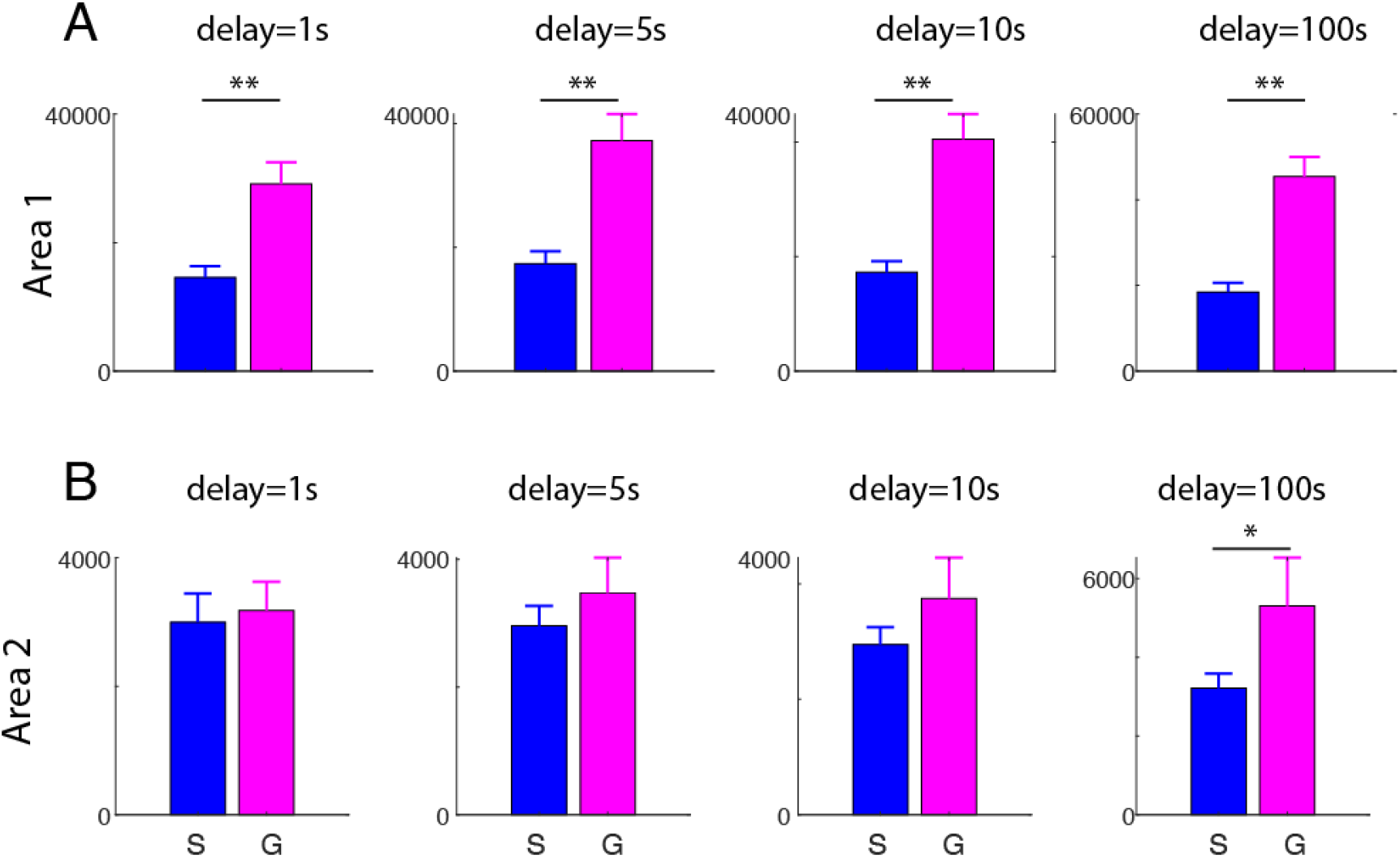
Group-housed mice have a larger speed transition area. Areas are calculated with boundaries at a low (10%) (**A**) and high (50%) (**B**) threshold. Group-housed mice show larger low threshold areas in all conditions (p<0.001 T-test).

### Parvalbumin-Cre mice do not show differences in running based on housing conditions

C57Bl/6 mice are commonly used as background for a variety of transgenic strains. A very common approach is to express Cre-recombinase under a cell type specific promoter in order to manipulate selective cell populations, e.g. neurons in different layers of the cerebral cortex (Daigle et al., 2018). We thus wondered if this genetic manipulation to express Cre could lead to subtle changes in running behavior. In order to investigate this question, we repeated our experiments with parvalbumin-Cre (PV-Cre) mice. Therefore, we measured wheel running in single-housed (n=12 cages, 6 female, 6 male) and group-housed PV-Cre animals (n=8 cages with n=3 animals each; 4 cages females, 4 cages males). We find that single-housed PV-Cre mice showed single peaks in their running speed distributions (Fig. 7A, B), but that these distributions did not differ from group-housed PV-Cre mice (Fig. 7A, B). In addition, the numbers of speed peaks did not differ between single- and group-housed PV-Cre mice. Moreover, analyzing speed transitions did not show a difference in the number of speed transition areas between single- and group-housed PV-Cre mice (Fig.8). However, the area ratio was larger for short delays, indicating short-term changes in running speed in group-housed PV-Cre mice.

**Figure 7:**
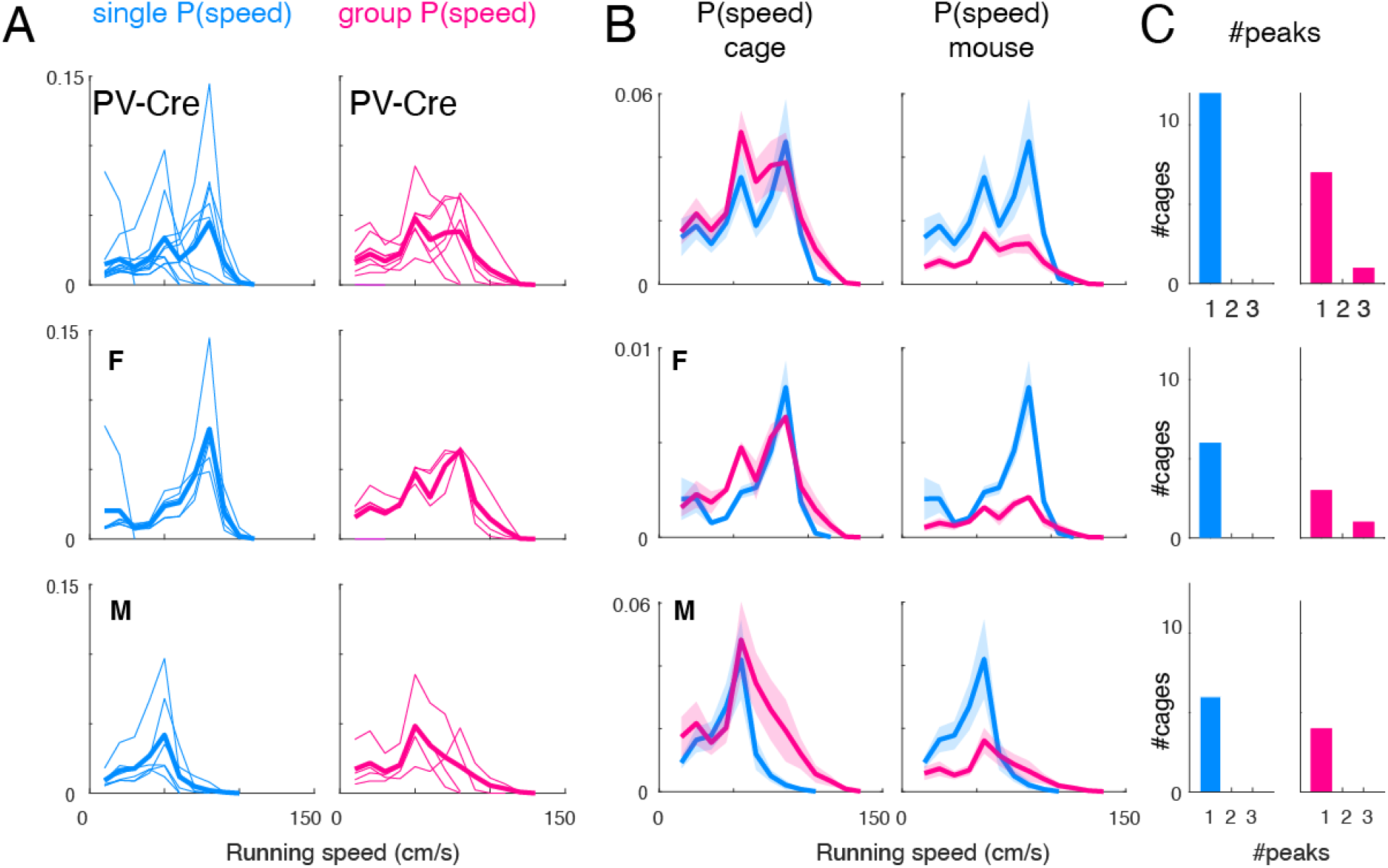
Running speed probability distributions of single-housed and group-housed PV-Cre mice. **A**: Running speed distributions for all cages (thin lines) and mean of distributions (thick line). **B**: Means and SEM of distributions in A per cage (left) and per mouse (right). **C**: Histogram shows number of peaks in distribution.

**Figure 8:**
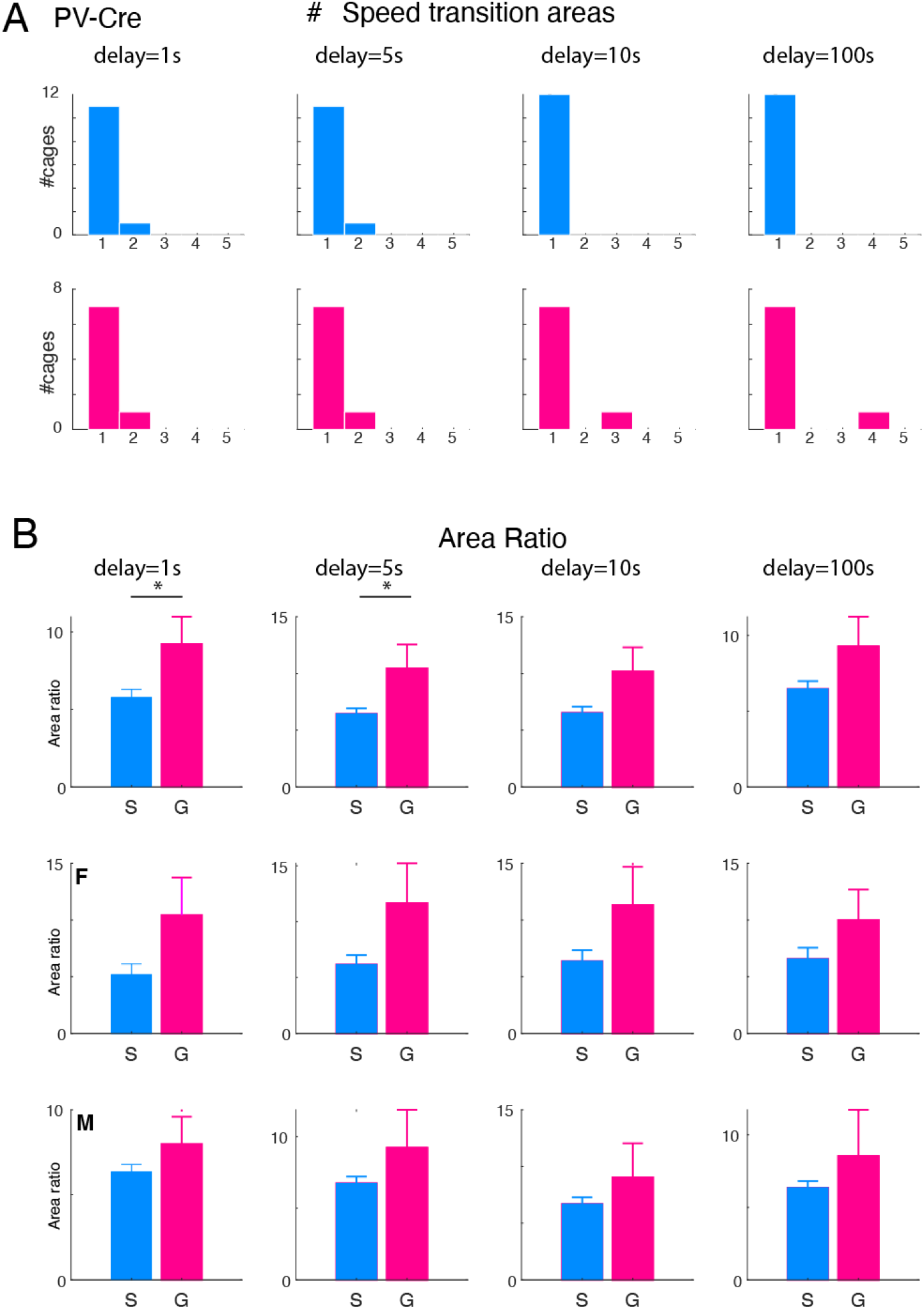
Running speed transitions of single-housed and group-housed PV-Cre mice. **A**: Histograms show the number of speed transition areas in single and group-housed mice for delays of 1s, 5s, 10s, 100s (all p>0.05 KS-test). **B**: Histograms show the area ratio in single and group-housed mice for delays of 1s, 5s, 10s, 100s. Lumped group-housed mice show a higher area ratio for short delays (p<0.05 T-test).

## Discussion

We used an instrumented running wheel to investigate the detailed running behavior of single-housed and group-housed mice. We find that group housing leads to a larger variability in running speeds and that this difference between single and group housing is most pronounced in female mice.

We find that single-housed females covered larger distances than males. We also find that group-housed mice contain the same mutual peak at a high speed (~80cm/s) as single-housed mice, but also contain one to two additional peaks at lower speeds than single-housed mice, meaning group-housed mice are more probable to spend more time running at the lower speeds than single-housed mice. We observed that single-housed female mice preferred to run at a narrow range of high speeds while running speeds of single-housed male mice were lower and were more variable. In contrast, in group-housed conditions, we find that male and female mice alike seem to run at multiple preferred speeds leading to multiple distinct peaks in the speed probability distribution. Since we group housed 3 mice and observed up to 3 peaks, these distinct peaks might indicate that individual mice ran at different preferred speeds. Alternatively, it could be that non-running mice might interfere with the running mouse, e.g., slowing down the wheel. Future work using video or RFID tagging is needed to discern this possibility.

In line with the observation of single peaks in the speed distributions, we find that speed transitions in single-house mice are limited to an almost binary switching between not running and running at a high characteristic speed for each mouse. For single-housed mice, we typically observed a single tightly circumscribed area indicative of stable running speed. This is consistent with prior observations suggesting that individual mice had preferred “cruising speeds” (De Bono et al., 2006). In contrast, in group-housed animals we observed that running speeds could vary both in the short-term (e.g., 1s) and the long-term (e.g., 100s). This is evidenced by the speed transition areas that were spread out over a larger area and could show multiple discrete areas indicating distinct combinations of speeds that were common. Thus, group-housed mice have a more variable running pattern in that they show a larger range of speed transitions.

This variability in running speeds in group-housed mice is consistent with the observation of multiple peaks in the speed distribution. Moreover, since mice transitioned from one non-zero running speed to another non-zero running speed, without stopping, this indicated that individual mice ran at different speeds during the running bout and thus were able to run at different graded cruising speeds. Thus, running behavior of individual mice seems to be more variable in group-housing conditions and thus their “cruising speed” might be set by a variety of social variables. Moreover, while multiple peaks in the running speed distribution could reflect the intrinsic speed preferences of individual mice housed together, the existence of the broad speed transition areas suggests that more variable running patterns of individual mice also contribute.

We observed that PV Cre single- and group-housed mice show fewer differences in running speed distributions and running speed transitions. This suggests that PV-Cre mice might have decreased behavior competition over the wheel or different group dynamics. These results indicate that expressing Cre in PV cells on a C57Bl/6 background can lead to subtle differences in running behavior from C57Bl/6 wild type.

In summary, when mouse locomotion was observed in detail in a variety of different housing conditions, with sex differences, and different genetic strains, distinct differences in running behavior due to group housing were observed. Our results suggest that our system is able to detect subtle changes in running behavior and could be used to detect changes in a variety of other models of neurological disorders.

## Materials and Methods

All animal procedures were approved by the NIH Animal Care and Use Committee. We used male and female adult mice (C57BL/6J background, Jackson Labs, #000664 and Parvalbumin-Cre mice Jackson Labs, #008069 backcrossed in-house on C57Bl/6J background).

We fitted a running wheel (Techniplast, USA) with neodymium permanent magnets (6×3 mm; e.g. Diymag). The magnets passed by a magnetic induction switch (Reed; contact normally open; plastic; Wowoone; 2.5mm×14mm). We acquired the switch opening/closing events with a National Instruments data acquisition board (NI USB-6215) at a 100Hz sampling rate using custom scripts in MATLAB (Mathworks).

Mice were group-housed and separated by sex for several weeks with access to a single wheel in each cage. Thus, they were well acclimated to the wheel before the start of the experiment (Bowen et al., 2016). At the chosen recording day, group running was assessed in their respective home cages. For single-housed animals, animals were single-housed for up to 72 hr with non-competitive access to a cage wheel attached to the data acquisition system. Locomotion activity for both group-housed and single-housed animals was continuously recorded at a sampling rate of 100 Hz/cage wheel for up 72 hr under a 12h:12h light-dark cycle.

Running activity was analyzed with custom-written code in MATLAB, and data is expressed as mean ± SEM if not stated otherwise. Means are compared by Student’s T-test, and distributions are compared by Kolmogorov-Smirnov (KS) tests. The peaks in the running speed distributions were detected using the ‘findpeaks’ function in MATLAB with ‘MinPeakProminanc’e set to 10%. The 2D running speed transition distribution areas were analyzed by using the ‘bwboundaries’ function in MATLAB (with the ‘’noholes’ option set) with thresholds of 10% and 50% of the maximum.

## Acknowledgments

DP designed and supervised the study. ZKT contributed to system design, system construction, and study. ZKT and DP analyzed data and wrote the manuscript. We thank Amber Tietgens for help during data acquisition. This research was supported by the Division of the Intramural Research Program (DIRP) of the National Institute of Mental Health (NIMH), USA, ZIAMH002797, ZIAMH002971 and the BRAIN initiative Grant U19 NS107464-01.

